# Metabolism constrains bird and mammal ranges and predicts shifts in response to climate change

**DOI:** 10.1101/167973

**Authors:** Lauren B. Buckley, Imran Khaliq, David L. Swanson, Christian Hof

## Abstract

**Aim:** We test whether physiological constraints on maximum metabolic rate and the factor by which endotherms can elevate their metabolism (metabolic expansibility) govern cold range limits for mammal and bird species.

**Location:** Global

**Methods:** We examine metabolic expansibility at the cold range boundary (ME_crb_) and its trait predictors and then use ME_crb_ to project range shifts for 210 mammal and 61 bird species.

**Results:** We find evidence for metabolic constraints: the distributions of metabolic expansibility at the cold range boundary peak at similar values for birds (2.7) and mammals (3.2). The right skewed distributions suggest some species have adapted to elevate or evade metabolic constraints. Mammals exhibit greater skew than birds, consistent with their diverse thermoregulatory adaptations and behaviors. Mammal and bird species that are small and occupy low trophic levels exhibit high levels of ME_crb_. Mammals with high ME_crb_ tend to hibernate or use torpor. Predicted metabolic rates at the cold range boundaries represent large energetic expenditures (>50% of maximum metabolic rates). We project species to shift their cold range boundaries poleward by an average of 3.9° latitude by 2070.

**Main conclusions:** Our analysis suggests that metabolic constraints provide a viable mechanism for projecting cold range boundaries and range shifts in response to climate change for endotherms.

## INTRODUCTION

Environmental temperatures govern the performance and energy use, and ultimately the abundance and distribution, of animals (Bozinovic *et al*., 2011). Performance and energetic constraints provide a powerful basis for projecting responses to climate change because the constraints should extrapolate better into novel environments than statistical correlations (Radeloff *et al*., 2015). Models translating environmental conditions into the body temperatures of ectotherms and quantifying limitations on performance and activity durations can robustly predict patterns of abundance and distribution (Kearney & Porter, 2009; Buckley *et al*., 2010). The translation is more complex for endothermic animals because they can use endogenous heat production to maintain their body temperatures under a wide range of environmental thermal conditions if available resources and physiological capacities are sufficient (Boyles *et al*., 2011; McNab, 2012). Thus, few mechanistic approaches predict endotherm distributions (but see examples reviewed in Boyles *et al*., 2011). Several recent examples employ biophysical models to estimate metabolic constraints, activity limitations, and water balance for focal endotherms (Kearney *et al*., 2016; Mathewson *et al*., 2017).

Fundamental physiological constraints on metabolism limit maximum metabolic rate and the factor by which endotherms can elevate their metabolism (Humphries *et al*., 2004; Stager *et al*., 2015). An initial test of metabolic constraints (Root, 1988) suggested that the cold range boundaries of passerine birds in North America coincided with winter metabolic rates at the cold range boundary being elevated by a factor of 2.5 over basal rates, but subsequent analyses (Repasky, 1991; Canterbury, 2002) have questioned the generality of metabolic constraints due to the limited biological, distributional, and environmental data available or poor fit between range boundaries and temperature isotherms. Physiological measurements indicate metabolic constraints and adaptations. Seasonal cold associated with either climate variability [Climate Variability Hypothesis (Janzen, 1967; Stevens, 1989; Ghalambor *et al*., 2006)] or seasonal temperature extremes [Cold Adaptation Hypothesis (Swanson & Garland Jr, 2009)] is expected to select for increased metabolic capacity. Maximum cold-induced metabolic rate (M_sum_) is greater in cold environments (Wiersma *et al*., 2007) and is phylogenetically conserved (Swanson & Garland Jr, 2009; Stager *et al*., 2015). Extensions of classic work examining how animals adapt to regulate heat (Scholander *et al*., 1950; Scholander, 1955) find that adaptation to environmental conditions alters both basal metabolic rate (BMR) and heat conductance in birds and mammals (Fristoe *et al*., 2015). Birds and mammals with more poleward range limits that experience colder minimum temperatures can tolerate colder temperatures without elevating metabolism (Khaliq *et al*., 2015).

Here we leverage extensive metabolic, distribution, and phylogenetic datasets (Khaliq *et al*., 2014; Fristoe *et al*., 2015) to test the viability of using metabolic constraints to project bird and mammal distributions. Specifically, we estimate the factor by which metabolism is elevated at the cold range boundaries (metabolic expansibility, ME_crb_). We expect the distribution of ME_crb_ to be normal and strongly peaked if the cold range edges of birds and mammals are limited by the capacity of their metabolic systems to maintain approximate temperature homeostasis. A peaked distribution would indicate similar limits to ME_crb_ across birds and mammals that differ substantially in geographic distribution, habitat, traits, and life history. However, positive skew in the distribution could reflect species that are metabolically adapted to or able to evade cold conditions. Species may evade extreme temperatures by adjusting activity times (e.g., diurnality) or the maintenance of body temperatures (e.g., use of hibernation or torpor) or by selecting favorable microclimates. We hypothesize that because mammals use strategies to evade full exposure to winter cold (e.g., hibernation, use of subnivean space) to a much greater degree than birds (Swanson, 2010; Williams *et al*., 2014; Ruf & Geiser, 2015), that mammals will show both greater skew in the density distribution of ME_crb_ and more values of ME_crb_/Msum approaching or exceeding unity than birds.

We test whether physiological, behavioral and ecological traits (body size, nocturnality, torpor use, diet) associated with adaptation or evasion correspond to higher ME_crb_ values. Body size influences the ability to use potential microclimates as well as metabolic rates and thermal inertia. Trophic levels influence the seasonal availability of food and metabolic rate (McNab, 2008, 2009). We also examine the conservatism of traits and metabolic expansibility at the cold range boundary (ME_crb_) across the phylogeny. Finally, evidence for metabolic constraints suggests that (in the absence of adaptation or acclimation) species will follow thermal isoclines through climate change. We thus project ranges and range shifts in response to predicted climate change.

## MATERIALS AND METHODS

We estimated the factor by which metabolism is elevated at the cold range boundaries (as in Root, 1988). We calculated metabolic rate (ml O_2_ h^−1^) at the cold range boundary as MR_crb_ = (T_lc_ − T_min_)C + BMR, where T_lc_ is the lower critical temperatures bounding the lower limit of the thermal neutral zone (TNZ); T_min_ is the coldest environmental temperatures (see below) at the cold range boundary; BMR is basal metabolic rate (ml O_2_ h^−1^), and C is thermal conductance (ml O_2_ h^−1^ °C^−1^) (figure 1). We calculated metabolic expansibility at the cold range boundary as ME_crb_=MR_crb_ /BMR. We focus on cold range boundaries because they are more likely governed by metabolic constraints than are warm range boundaries (see discussion); at warm range boundaries the capacity for evaporative cooling may be more limiting than the associated metabolic costs and minimal endogenous heating is favored (Tieleman & Williams, 2000; McKechnie *et al*., 2016b). Our estimates of ME_crb_ are approximate in that they do not account for additional factors such as use of solar radiation, convective heat loss, microclimate variation, and microhabitat selection (Porter & Kearney, 2009).

**Figure 1.**
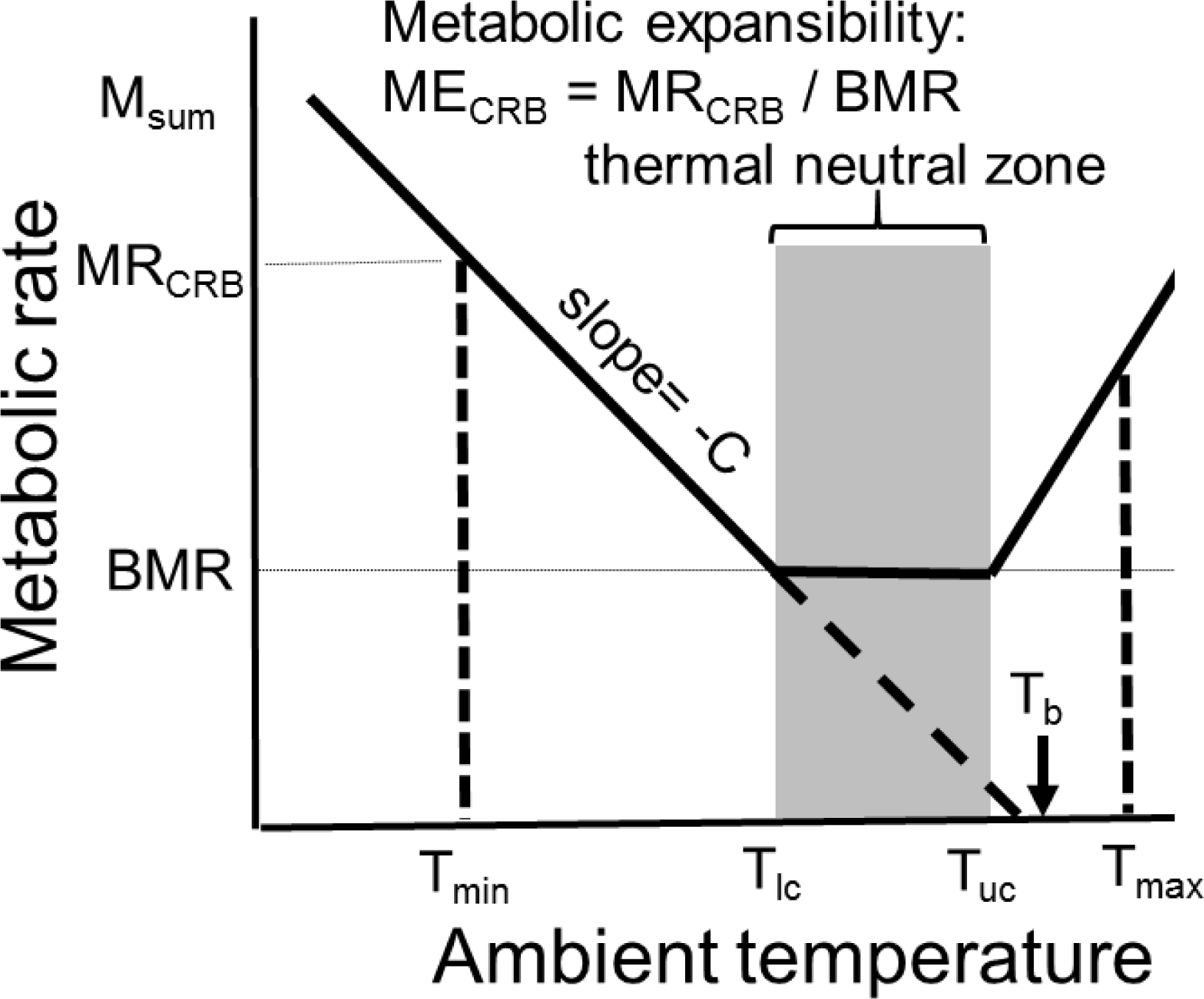
How ambient temperature governs metabolic rate. The thermal neutral zone [bounded by lower (T_lc_) and upper (T_uc_) critical temperatures] is the range of temperatures over which endotherms are able to maintain their basal metabolic rate (BMR). We use the minimum (T_min_) and maximum (T_max_) ambient temperatures across a species’ range to estimate sustained metabolic rate at the range boundary (MR_crb_). We calculate metabolic expansibility (ME_crb_) as MR_crb_/BMR and depict maximum metabolic capacity (M_sum_). Thermal conductance (C) is calculated as the slope of the line terminating at body temperature (T_b_).

### Data

We restricted our analysis to resident (non-migratory) species. We omitted species on islands and those with latitudinal range limits constrained by continental boundaries. We additionally restricted our analysis to cold range boundaries with temperatures below the T_lc_ (omitted 1% of species). Accounting for these constraints and limitations on available physiological data, we analyzed 210 and 61 cold range boundaries for mammal and bird species, respectively.

For species distribution data, we used the IUCN range maps for mammals (Patterson *et al*., 2005) and the BirdLife range maps for birds (BirdLife International and NatureServe, 2014). We calculated temperatures at the range boundaries (T_min_ and T_max_) using BIO5 (max temperature of warmest month) and BIO6 (min temperature of coldest month) at 5 minute resolution from the WorldClim dataset (Hijmans *et al*., 2005). Data are interpolated from air temperature at weather stations (generally 2m high). Trait data are insufficient to account for microhabitat use (e.g., burrows or under snow), but our trait analysis does provide some indication of exposure to air temperatures. We extracted the grid cells at the northern and southern extremes of the species’ distribution for each 5-minute longitudinal band. We quantified the degree to which range boundaries follow temperature isoclines as the standard deviation and median absolute deviation (mad, R function mad) of cells along the range boundaries. Subsequently, we estimated T_min_ and T_max_ as the median of the grid cells along the cold and warm range boundaries, respectively. We checked that minimum and maximum temperatures were sufficiently constant across the range boundaries for our results to be robust to our selection of the median (figure S1). Current data are normals for 1950-2000 and future data are downscaled global climate model (GCM) projections from CMIP5 (IPCC Fifth Assessment) averaged over 2061-2080. We examined output from both the HadGEM2-AO and CCSM4 models assuming a midrange greenhouse gas concentration scenario (Representative Concentration Pathway RCP6.0, indicates a 6W/m^2^ increase in radiative forcing in 2100 relative to pre-industrial values, http://cmip-pcmdi.llnl.gov/cmip5/). Data were accessed using the getData function in the R package raster.

The bounds of the TNZ (T_lc_ and T_uc_, °C) and body temperature (T_b_, °C), were compiled from the literature by Khaliq et al. (2014). We incorporated data compiled for additional species (Canterbury, 2002; Riek & Geiser, 2013; Bozinovic *et al*., 2014). We used BMR data from Fristoe et al. (2015) and McNab (2008, 2009) after assessing whether the data met criteria for data quality (see below). We extracted Msum data (Msum = BMR + maximum thermogenic capacity) for 20 mammal and 6 bird species from existing compilations (Rezende *et al*., 2004; Lovegrove, 2005; Swanson & Garland Jr, 2009; Stager *et al*., 2015). Those MR values reported in watts were converted to oxygen consumption assuming a factor of 179 ml O_2_ h^−1^ W^−1^, which corresponds to lipid metabolism (Schmidt-Nielsen, 1997). Minimum conductance was estimated as the absolute value of the slope of the line connecting T_lc_ at BMR to T_b_ when metabolic rate is 0: Cmin=|(0-BMR)/(T_b_-T_lc_)|(Fristoe *et al*., 2015). We use units of oxygen consumption for metabolism and conductance to align with previous analyses (Fristoe *et al*., 2015).

Analysis of the quality of the data compiled in Khaliq et al. (2014) identified issues with T_uc_ but not T_lc_ (McKechnie *et al*., 2016a). Some Tuc data are of lesser quality due to small sample sizes or weak measurement protocols (McKechnie *et al*., 2016a), so we only use the T_uc_ data for a coarse analysis mentioned in our discussion. We omitted T_uc_ measurements that were found to be of poor quality [“No UCT” or “NA-” categories; we kept values based on low sample sizes due to the tentative nature of our analyses] (McKechnie *et al*., 2016a). We revisited the source papers to assess whether the T_lc_ data were calculated from valid BMR measurements. We used the following criteria to assess data quality for BMR: measurements were made during the rest phase on inactive individuals in a postabsorptive state. We additionally recorded whether individuals measured were field-collected (or the first generation reared in a laboratory or zoo in a small number of cases) and the location of field collection, as individuals collected far from the range boundary may lack adaptations and acclimation present near the range boundary. We assessed the influence of these quality criteria on our results as described below.

Diet, habitat, and nocturnality data were extracted from Elton Traits (Wilman *et al*., 2014). Data on whether a species uses torpor or hibernation were extracted from McNab (2008, 2009) and Ruf and Geiser (2015). A “torpor” trait was assigned a value of 1 if the species uses either torpor or hibernation and 0 otherwise. Data on relevant thermoregulatory traits such as body shape, insulation, and fur or feather properties were inadequate to include the traits in the analysis.

### Analyses

We examine the distribution of ME_crb_ estimates across birds and mammals to assess evidence for a metabolic constraint. We assessed skewness and kurtosis of the ME_crb_ distribution using the skewness metric and D’Agnostino skewness test and Geary metric and Bonett-Seier test in the R moments package. We tested for unimodality in the distributions using Hartigans’ dip statistic in the R diptest package. To test whether ME_crb_ varies systematically with T_min_ or T_max_, we constructed null models for ME_crb_ by randomizing T_min_ or T_max_ among species and calculating the median and mean ME_crb_ values. We repeated the randomization 1000 times.

We then used regressions to assess whether species’ traits indicating adaptation to or evasion of cold temperatures can explain variation in ME_crb_. We used model selection based on AICc (dredge function) and model averaging (model.avg function in R package MuMIn) to conclude that the best models omitted interactions between the predictor variables (mass, diet, nocturnality, torpor). Accounting for phylogeny did not alter our results, so we report phylogenetic analyses in appendix S1.

We used thermal isoclines (consistent with species maintaining a constant ME_crb_ in the absence of acclimation or adaptation) to project species’ distribution in both current and future environments. We restricted predicted distributions to observed west and east longitudinal extents since we have no basis for predicting longitudinal distributions. We identified all pixels with T_min_ warmer than the predicted physiological lower temperature limit (based on species-specific observed ME_crb_). We removed pixels predicted to be thermally habitable that were geographically isolated from other habitable pixels from our prediction using the clump function in the R package raster. We omitted all clumps with areas less than 5% of the area of the largest clump, because the core of the predicted distribution is most representative of latitudinal extents. We further restricted our predicted distribution to clumps overlapping with the latitudinal extent of the observed species range. We then quantified the median latitude of grid cells along the cold range edges.

## RESULTS

### Metabolic expansibility at the cold range boundary

Our analysis of bird and mammal species with disparate geographic distributions, habitats, traits, and life histories suggest that winter temperatures and the ability to elevate metabolism to maintain body temperatures constrain cold range boundaries (Fig. 2). Cold range boundaries of both mammals (sd=4.6°, mad=3.9° of median T_min_) and birds (sd=4.6°, mad=3.3° of median T_min_) approximately follow temperature isoclines (Fig. S1 in the Supporting Information). Translating this thermal variability into metabolic consequences, the median standard deviations represent a change in metabolic rate of 12.4±12.2% (mean ± sd) for mammals and 10.8±7.8% for birds. The distributions of metabolic expansibility, ME_crb_, are peaked and peaks occur at similar values for birds and mammals. The bird distribution has a slight dip at the peak of the density distribution, which we attribute to limited sample size in the absence of evidence for non-unimodality (Hartigans’ dip test: D=0.05, p=0.3). We thus estimate the peak value as the mean of the two subpeaks. The density distribution of ME_crb_ peaks at 2.72 for birds (median=3.21, mean ± sd= 3.28 ± 1.63) and at a slightly higher value (3.17, median=3.63, mean ± sd= 4.64 ± 3.35) for mammals. ME_crb_ values fall outside the 95% confidence intervals of the null model estimated by randomization for both mammals (median: 3.53-3.54, mean:4.55-4.56) and birds (median: 2.63-2.64, mean:3.16-3.17). The previous value found for birds (2.5 × BMR; Root, 1988) was similar to our estimate of the peak of the distribution.

**Figure 2.**
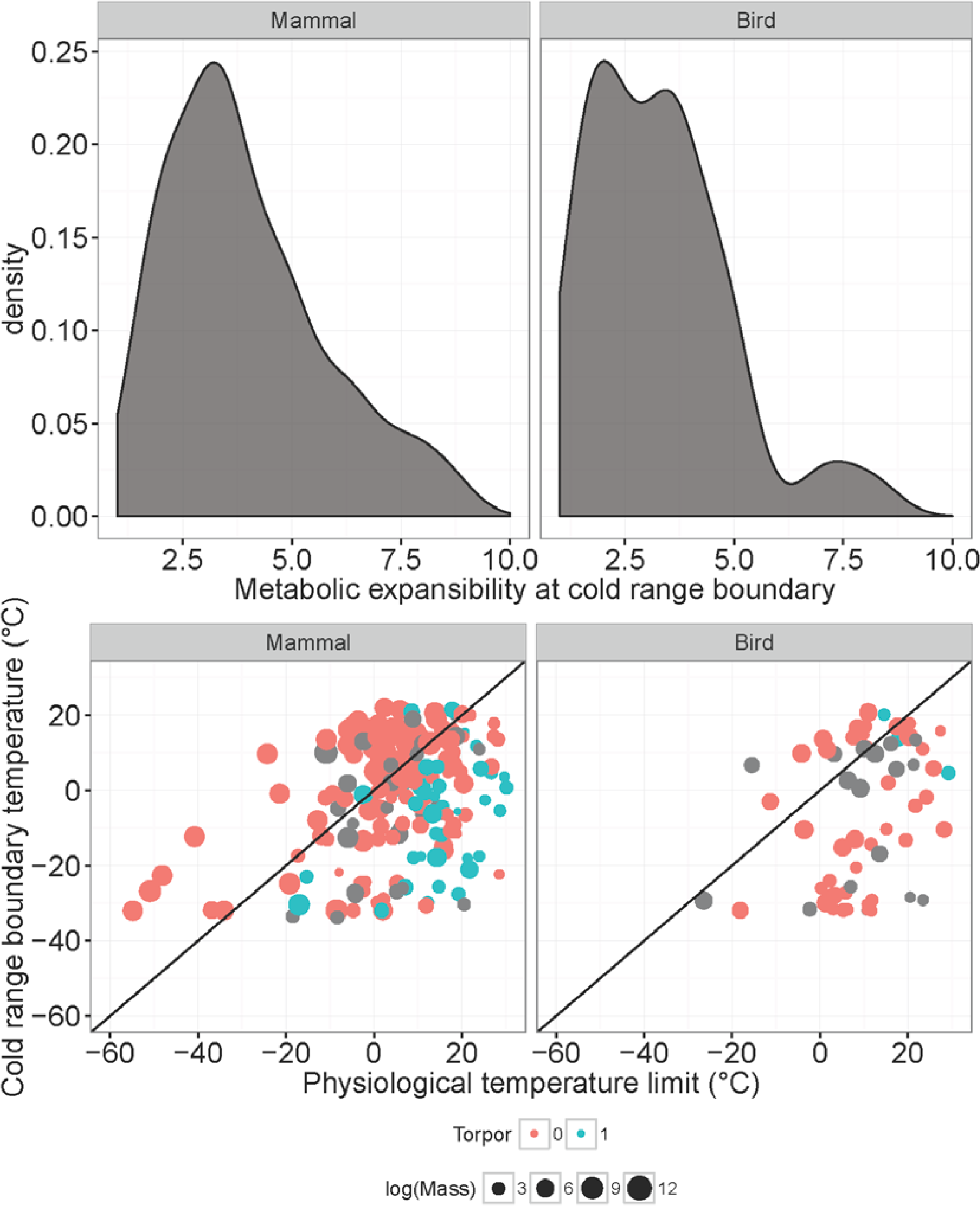
The density distribution of metabolic expansibility, ME_crb_ (the factor by which metabolic rate at the cold range edge is elevated over basal metabolic rate) peaks at similar values for birds and mammals (a). We examine interspecific variation in ME_crb_ by (b) plotting the physiological temperature limit predicted by assuming the mode of ME_crb_ and the observed temperatures at the cold range boundaries. Mammals and birds that are small (symbol size) and use torpor or hibernation (color, 1=use, gray=no data) tend to be found in environments colder than predicted assuming the mode ME_crb_ (i.e., they have higher ME_crb_).

We assessed whether ranges may be constrained more strongly by maximum metabolic capacity (M_sum_) rather than the factorial capacity for elevating metabolism over BMR (ME_crb_). Among the limited data available for our focal species (N=20 mammal and 6 bird species), M_sum_ is on average 5.0 times BMR (median 5.4, 25^th^ to 75^th^ percentile= 4.0 to 6.3). The density distribution of the ratio of MR_crb_ to M_sum_ peaks at 0.7 (median=0.88, mean + sd =0.96 + 0.44, Fig. 3).

**Figure 3.**
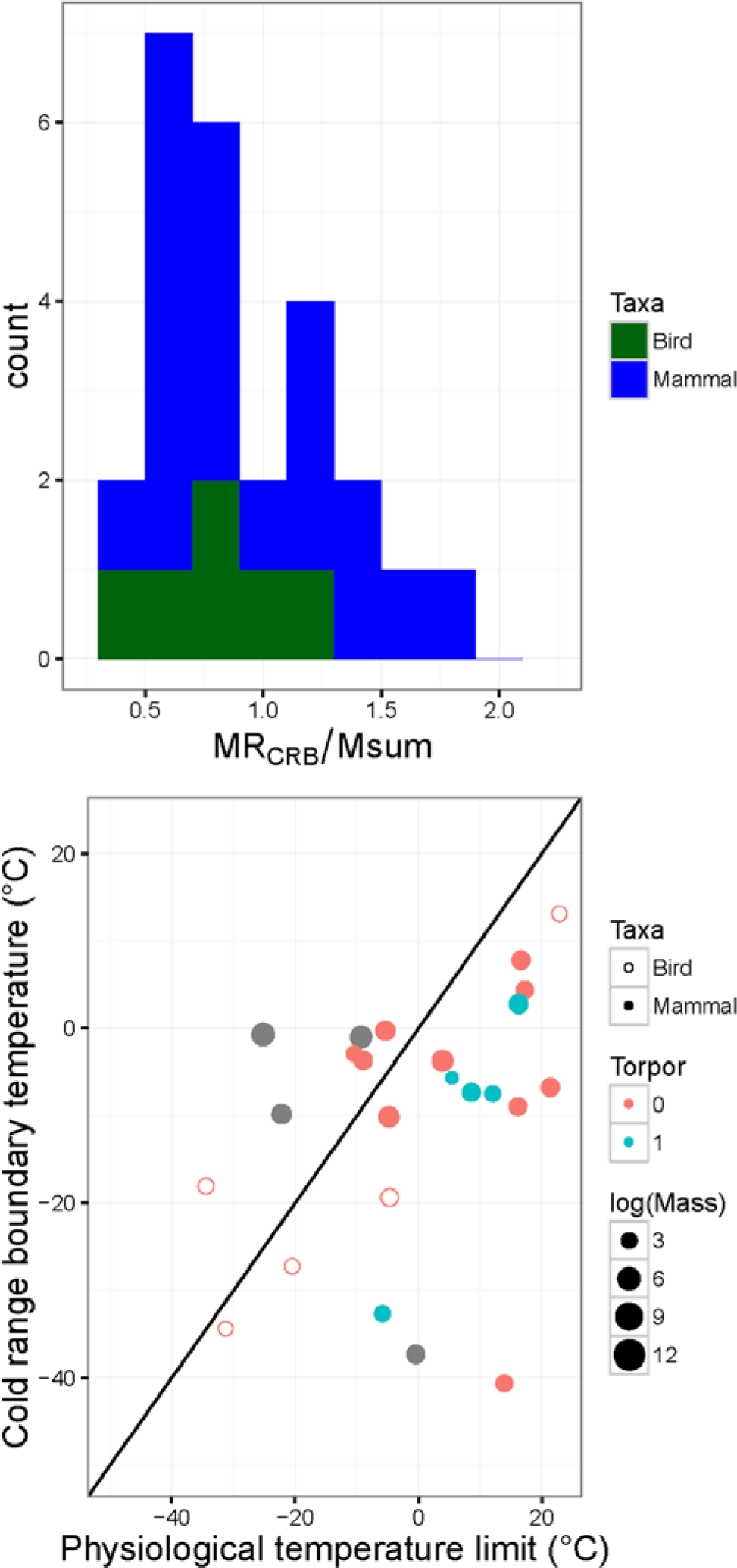
A histogram (a) of the ratio of maximum metabolic capacity (M_sum_) to estimated metabolic rate at the cold range boundary (MR_crb_) suggests the high energetic demands of thermoregulation. We examine interspecific variation in the ratio (MR_crb_ / M_sum_) by plotting the observed temperatures at the cold range boundaries against the physiological temperature limit corresponding to MR_crb_ = 0.7 M_sum_ (b). We depict mammals (filled circles) and birds (hollow circles), mass (symbol size), and use of torpor or hibernation (color, 1 indicates use).

The right skewed distributions of ME_crb_ (Fig. 2) suggest that some species have evolved the capacity to maintain a higher ME_crb_ or to evade the constraints of cold temperatures via torpor, microclimate selection, or movement. The distribution for mammals is more skewed (2.54) than that for birds (1.06), but both exhibit significant positive skew (D’Agnostino test, mammals: z=9.52, p<10-^15^; birds: z=3.22, p<0.001). Only mammals exhibit significantly more kurtosis than expected under normality (Bonett-Seier test, mammals: Geary metric: 0.78, z=10.20, p<10-^15^; birds: Geary metric: 0.66, z=0.78, p=0.2).

We next assess whether traits that allow organisms to maintain high metabolism or evade cold temperature can explain the skewed distribution. Mammalian traits (mass, diet, nocturnality, and use of torpor or hibernation) account for a substantial portion of variation in ME_crb_ (r^2^=0.29, F_[7,171]_=11.2, p<0.001); mammal species that are relatively small (t=−4.36, p<0.001) and use torpor or hibernation (t=4.99, p<0.001) tend to have higher ME_crb_ (Fig. 2, Table S1). Diet also significantly influences ME_crb_, with granivores having higher ME_crb_ than mammals consuming other diets (F=2.44, p<0.05, ANOVA, Table S3). Bird traits (mass, diet, and nocturnality) likewise account for a substantial portion of variation in ME_crb_ (r^2^=0.28, F_[6,54]_=4.9, p<0.001); birds that are small (t=−2.82, p<0.05) tend to have higher ME_crb_. Birds that eat invertebrates or plants and seeds exhibit higher ME_crb_ than those consuming other diets (F=7.85, p<0.01, ANOVA). Limited phylogenetic signal in mammal and bird ME_crb_ (Fig. S3) arises largely from conservatism of predictor traits (appendix S1). Phylogenetic regressions do not substantially deviate from linear regressions (Table S1, appendix S1).

### Data quality

In our full dataset for mammal ME_crb_, the following proportions of species with data met our BMR quality control criteria: 93.9% [58.6% including NA (not available) values as not meeting quality criteria] were measured during the resting phase, 70.0% (42.9% including NA values) were postabsorptive, and 81.2% were wild-caught (70.0% including NA values) (Table S2). Of the quality criteria, only whether the mammal species was live-trapped or captive was a significant predictor of ME_crb_ (resting phase: F_[1,82]_=1.53, p=0.22; postabsorptive: F_[1,82]_=0.04 p=0.84; wild caught: F_[1,82]_=5.15, p<0.05). However, restricting the dataset to wild-caught species does not substantially alter the peak value of metabolic expansibility (peak= 3.27, mean= 4.72, median= 3.75). The trait predictors of ME_crb_ remain similar when considering only wild-caught individuals (Table S3).

In our full dataset for bird ME_crb_, the following proportions of species with data met our BMR quality criteria: 93.0% (86.9% including species without data) were measured during the resting phase, 88.9% (52.4% including species without data) were postabsorptive, and 68.9% were wild-caught (50.8% including species without data) (Table S2). Similar to mammals, of the quality criteria only whether the bird species was live-trapped or captive was a significant predictor of ME_crb_ (resting phase: F_[1,32]_=0.00, p=0.94; postabsorptive: F_[1,32]_=0.68 p=0.42; wild caught: F_[1,32]_=6.90, p<0.05). However, restricting the dataset to wild-caught species did not substantially alter the peak value of ME_crb_ (peak= 2.60, mean= 3.07, median= 3.11). The trait predictors of ME_crb_ remained similar, but some predictors lose significance, when considering only wild-caught individuals (Table S3).

The measured individuals were collected throughout the species’ range with average positions near the center of the range for both mammals (median and mean from range edge: 10.3° and 13.4° latitude, 47.2% and 49.4% of the species’ latitudinal range) and birds (median and mean from range edge: 18.3° and 20.4° latitude, 50.8% and 49.9% of the species’ latitudinal range). Neither distance metric is a significant predictor of ME_crb_ in mammals (distance: F_[1,142]_=0.27, p=0.60; percent: F_[1,142]_=0.10, p=0.76) or birds (distance: F_[1,29]_=0.12 p=0.73; percent: F_[1,29]_=0.39 p=0.54). Collection locations have a median elevation of 275m (25^th^ and 75^th^ quantiles: 39 to 849m, based on collection coordinates and Google Maps Elevation API). Thus, few of the physiological measurements reflect metabolic adaptation to high elevation.

### Range shifts

We forecast potential range shifts by examining how metabolic constraints will shift through climate change. For example, North American rodent species differ in their metabolic constraints, the extents of their current distribution, and the projected range expansion as a result of climate change (Fig. 4 for projections using the HadGEM2-AO model; Fig. S4 for CCSM4 model projections). The quality of the range projections vary across species (Figs. S5-S8). We predict that most mammals and birds will shift their cold range boundaries poleward through climate changes (Fig. 4). We project a similar magnitude of cold range boundary shifts for mammals (mean=3.77°, median=2.58°) and birds (mean=4.20°, median=3.63°). Numerous species are projected to shift their cold range boundary poleward by 6° latitude (75% quantile) and some species are predicted to shift by as much as 22° (Fig. 4).

**Figure 4.**
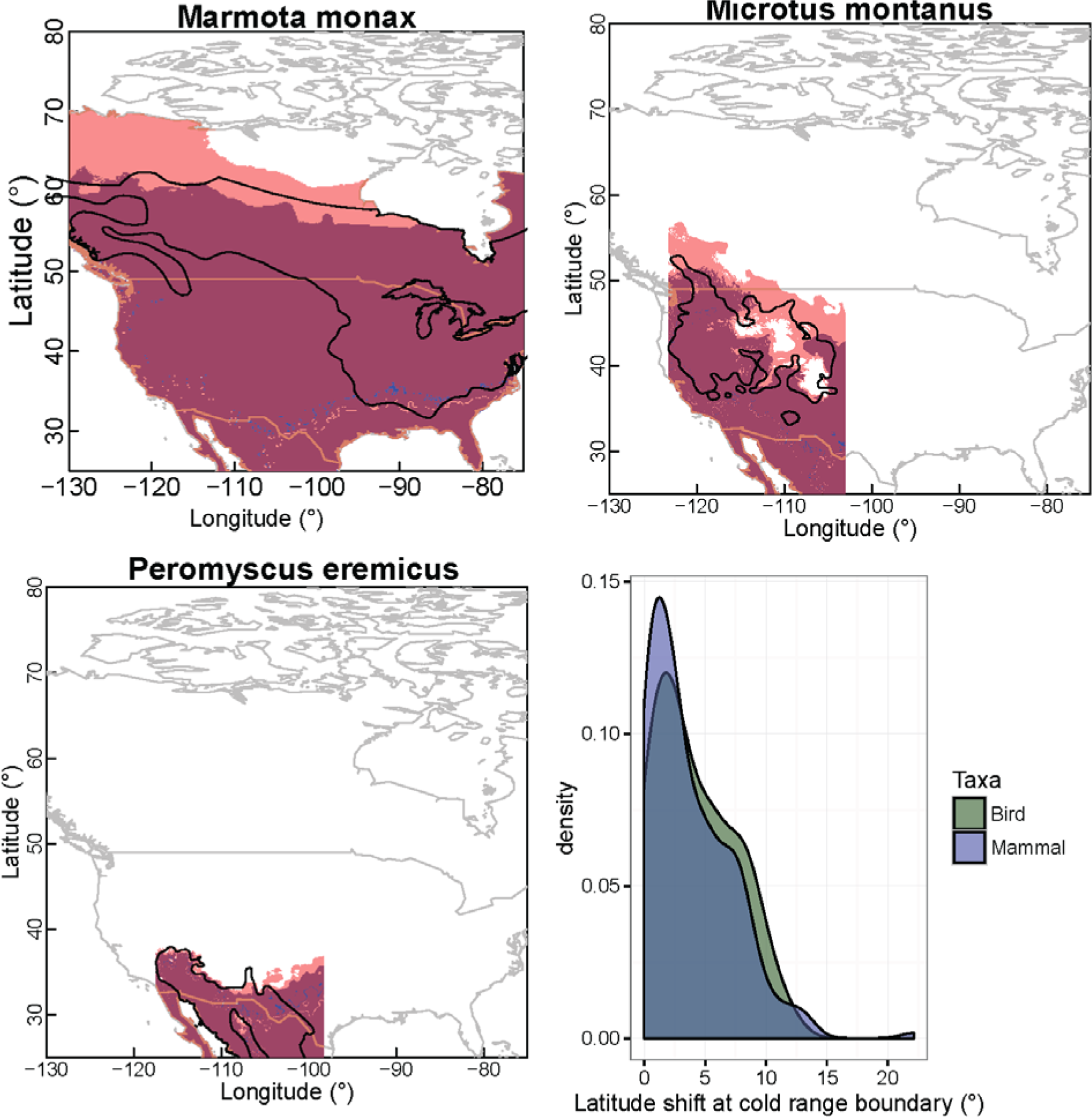
We depict observed cold range boundaries (black polygons: IUCN range maps) and those projected based on metabolic constraints for exemplar North American rodents in current (blue: 1950-2000) and predicted future (red: 2061-2080 from HadGEM2-AO model) climates (a – c). Purple shading indicates portions of the projected range occupancy that persists through climate warming. We note few areas of range contraction (blue). The species differ in the extent of their current distribution and the projected range expansion as a result of climate change (a, *Marmota monax*, groundhog; b, *Microtus montanus*, montane vole; and c, *Peromyscus eremicus*, cactus mouse). Projected based on metabolic constraints indicate that the majority of mammals (purple) and birds (green) will shift their cold range boundary modestly poleward through climate changes (d). However, numerous species are projected to shift their cold range boundary poleward by 10° latitude and some species are projected to shift by as much as 22°.

## DISCUSSION

Our analysis supports a mechanism underlying observations that endotherms track thermal isotherms through climate change (Tingley *et al*., 2009; Chen *et al*., 2011). However, many observed range shifts have been idiosyncratic in extent and direction (Gibson-Reinemer & Rahel, 2015). Filtering the range shifts through the lens of metabolic constraints may resolve some discrepancies. Our data are consistent with the poleward range edges of both birds and mammals being constrained by the factor by which they can elevate their metabolism above basal rates (perhaps resulting from a constraint on maximum metabolic rates). The constraint may result from either direct physiological limitations on metabolism, such as the ability to sustain high rates of thermogenesis over prolonged periods, or limitations on energy acquisition. The rates of ME_crb_ that we find for birds (peak of distribution=2.7) are similar to a previous value (2.5) for a more taxonomically and geographically restricted analysis (Root, 1988).

We find a slightly higher peak of the ME_crb_ distribution for mammals (3.2). The distribution of ME_crb_ is right skewed, more so for mammals than for birds. The greater skew in the mammal ME_crb_ distribution is consistent with the prominent use of hibernation and protected microclimates (e.g., burrows, dens, subnivean space) during winter in mammals, but lesser use of these options to avoid cold thermal environments in birds (Swanson, 2010; Ruf & Geiser, 2015). These adjustments have the effect of rendering the thermal conditions encountered at the ME_crb_ as less extreme than the actual ambient conditions, which results in an overestimation of the thermal isocline followed by the cold range boundary. In addition, differences in the mechanisms of thermoregulation between mammals and birds may contribute to the difference in ME. Cold-adapted mammals have well developed capacities for non-shivering thermogenesis through brown fat, but birds lack brown fat and although they may possess some muscular non-shivering thermogenesis, muscular shivering appears to be the primary mechanism of heat production in birds (Mezentseva *et al*., 2008).

The limited data on maximum cold-induced metabolic capacity (M_sum_) provide additional support for a metabolic constraint. We estimate that thermoregulation at the cold range boundary requires a substantial proportion (>50%) of the potential metabolic capacity for thermogenesis of the species. This supports the existence of a metabolic constraint on range boundaries and suggests that species use a substantial portion of their maximum metabolic capacity to thermoregulate. The right skewed distribution (and instances where MR_cb_/M_sum_ >1) suggests that some species use torpor or hibernation or evade the coldest temperatures through habitat and microclimate selection (Fig. 3). Because M_sum_ is a flexible trait correlated with environmental conditions (Rezende *et al*., 2004; Swanson, 2010), ratios approaching or exceeding one may also result from M_sum_ measurement occurring for populations in warmer climates than those at the cold range boundary. Correlations between M_sum_ and environmental temperatures have been previously documented for rodents (Rezende *et al*., 2004; Bozinovic *et al*., 2011) and birds (Swanson, 2010; Stager *et al*., 2015).

We identify traits associated with high values for ME_crb_, which may be adaptations to or consequences of inhabiting cold environments. Body mass is an important factor that influences ME_crb_. Smaller mammals, which tend to exhibit greater ME_crb_, may be able to evade cold temperatures through seeking shelters or selecting favorable microclimates. Alternatively, the ability to use torpor or hibernation enables mammals to inhabit colder environments. Mammals using torpor tend to be small, which may contribute to the relationship between mass and ME_crb_ (Ruf & Geiser, 2015). Small mammals may also be able to meet the resource requirements or store energy to maintain high metabolism through cold periods (due to the low per-organism, or total, metabolic rate stemming from their small size) (Humphries *et al*., 2004; Angilletta *et al*., 2010). Mammals at lower trophic levels (herbivores and invertebrate consumers) tend to exhibit higher ME_crb_. These species tend to have lower BMR (Khaliq *et al*., 2014, 2015) and their food sources may be more consistently available.

Lower mass-specific rates of heat production and heat loss (conductance) and smaller surface area to volume ratios favor larger body sizes in colder environments (i.e., Bergmann’s hypothesis, Ashton *et al*., 2000). Regardless, birds’ and mammals’ body sizes are diverse across climates (Khaliq *et al*., 2014; Fristoe *et al*., 2015). An analysis of regression residuals suggests that adaptations to cold environments in birds and mammals results in increased BMR and reduced conductance (Fristoe *et al*., 2015). Our analysis suggests that greater values of ME_crb_ (perhaps associated with selection for higher M_sum_) enable small birds and mammals to inhabit cooler environments. Birds from cold climates tend to exhibit higher M_sum_ (Stager *et al*., 2015). We identify traits (small body size, use of torpor or hibernation, diet) that may enable the elevated ME_crb_.

We omit analysis of warm range boundaries because we estimated that 61% and 45% of mammal and bird species with unconstrained warm range boundaries, respectively, do not experience T_max_ values exceeding their T_uc_. We note that these values are likely an overestimate because they do not account for heat associated with solar radiation or heat extremes, but they do suggest a greater viability for using metabolic constraints to project cold range boundaries. Our estimates of metabolic expansibility at the warm range boundry (for species with T_max_>T_uc_, following methodology for ME_crb_) approximate 1, highlighting the physiological challenges of heat dissipation (Weathers, 1981). At warm range boundaries, the capacity for evaporative cooling may be more limiting than the associated metabolic costs and minimal endogenous heating is favored (Tieleman & Williams, 2000; McKechnie *et al*., 2016b). Evaporative cooling poses a risk of dehydration in response to short term heat stress (McKechnie *et al*., 2012) and presents a challenge for longer term water balances (Kearney *et al*., 2016). Additionally, other biotic factors such as species interactions and resource or habitat constraints often constrain warm range boundaries (Sexton *et al*., 2009).

The observation that many species do not currently face metabolic constraints at their warm range boundaries suggests that direct temperature effects on metabolism resulting from climate change may predominately expand cold range boundaries rather than contract warm range boundaries. However, the increase in extreme heat events associated with climate change will likely result in range contractions via thermal stress (McKechnie & Wolf, 2009; Buckley & Huey, 2016). Range contractions at warm range boundaries may primarily result from indirect effects (e.g., species’ interactions), which often predominate in climate change responses (Tylianakis *et al*., 2008; Walther, 2010).

Assuming species follow thermal isoclines due to metabolic constraints, we project that species will shift their cold range boundaries poleward by an average of 3.9° latitude with numerous species shifting by 6° (75% quantile). Our analyses suggest that hibernation and torpor are important determinants of cold range boundaries. Climate change will also likely alter the energetics of hibernation, which may amplify poleward range shifts (Humphries *et al*., 2002). Many bird and mammal species rely on seasonal migration to obtain resources to meet seasonal energetic demands; considering the costs and benefits of such movements will be important to forecasting responses to climate change among migratory birds and mammals (which we excluded from our analysis) (Robinson *et al*., 2009). Shifting activity times may also function to modify estimates of range shifts (Levy *et al*., 2012).

Our analysis of a taxonomically and geographically diverse dataset suggests that metabolic constraints provide a viable mechanism for projecting the poleward range boundaries of endotherms. We hold that the Scholander-Irving model we employ provides a useful approximation of metabolism, but highlight several considerations that may improve upon the analyses. We estimate metabolic costs assuming homeothermy, but accumulating data suggest that endotherms represent a continuum of heterothermy (Boyles *et al*., 2013). Consideration of the occurrence of torpor/hibernation in the present study likely only partially accounted for deviations from thermoregulation due to T_b_ variation. Many endotherms seasonally acclimatize their insulation, behavior, and physiology (Boyles *et al*., 2011; Bozinovic *et al*., 2011). Some metabolic estimates in our database are specific to the cold season, but we did not fully account for acclimatization. A comparison of BMR and field metabolic rates (FMR) for small mammals failed to find support for intrinsic limitations on metabolism and low FMRs in very cold climates indicated acclimatization including behavioral avoidance (Humphries *et al*., 2005). Over longer time periods, adaptation may alter morphology or metabolic constraints (Boyles *et al*., 2011). Behavioral strategies for buffering cold include sheltering, huddling, basking, and microclimate selection (Angilletta *et al*., 2010). Despite these complications, metabolic constraints provide an initial step towards generalizable and mechanistic projections of endotherm responses to climate change.

## Acknowledgements

We thank N. Bouzid, A. Cannistra, S. Graham, J. HilleRisLambers and R. Huey for constructive input; T. Root for an introduction to metabolic constraints; and those who collected, compiled, and disseminated the data we used. We thank A. McKechnie for suggestions on assessing data quality. This work was supported by the National Science Foundation [DBI-1349865 to L.B.B. and OIA-1632810 to D.L.S.].

## Data accessibility

Data and scripts will be available from a repository upon publication. Data and scripts for the analysis are currently available at github.com/lbuckley/tnz. Data are temporally uploaded as an appendix.

## Biosketch

The research team is interested in the broad scale implications of physiology. The team integrates expertise in bird physiology (Swanson), bird and mammal macroecology (Khaliq and Hof), and energetic modelling (Buckley).

## Supporting Information

Appendix S1. Phylogenetic analysis.

Appendix S2. Data description.

Appendix S3. Data.

Table S1. Regression model results.

Table S2. Summary of data quality.

Table S3. Regression model results for live trapped animals.

Figure S1. Comparison of range edge temperature metrics.

Figure S2. Metabolic expansibility at the warm range edge.

Figure S3. Phylogenetic conservatism of metabolic expansibility and mass.

Figure S4. Range projections using CCSM4 climate projections.

Figure S5-S8. Range projections for all species.

